# The physiological dynamic clamp allows insect flight muscle to transition between two actuation modes in virtual reality

**DOI:** 10.64898/2026.07.31.742123

**Authors:** Ethan S. Wold, Rundong Yang, Ellen Liu, Nick Gravish, Simon Sponberg

**Affiliations:** School of Biological Sciences and School of Physics, Atlanta, GA, 30332 USA; Mechanical and Aerospace Engineering, University of California San Diego, San Diego, CA 92161, USA; Georgia Institute of Technology, Atlanta, GA, 30332 USA; Molecular Genetics and Cell Biology, University of Chicago, Chicago, IL 60637, USA

## Abstract

In most muscles, contraction is initiated by neural activation. Some groups of insects break this rule, flapping at frequencies far exceeding the neural drive to their flight muscles. These insects’ muscles (termed asynchronous) produce force in response to stretch, enabling flight at faster frequencies than would be possible through the slow calcium-dependent processes associated with neural activation. The first flapping insects lacked stretch-activated physiology, which then evolved on top of neural activation dynamics before likely being reduced again in some groups including moths. Stretch and neural activation can co-exist, but it remains unclear if stretch-activation alone is sufficient to generate asynchronous flapping in insect flight muscle. Building on prior closed-loop muscle physiology platforms, we develop a new way to perform a gain-of-function muscle physiology experiment called the physiological dynamic clamp. Inspired by dynamic clamp experiments in neuroscience, we couple isolated intact flight muscle from a hawkmoth, *Manduca sexta*, to simulated stretch-activation in virtual reality. Tuning virtual reality parameters allows us to manipulate the degree of stretch-activation *in silico* while retaining all other physiological properties of the muscle. With artificially enhanced stretch activation, we find that hawkmoth muscle can support stretch-activated work at typical wingbeat frequencies. When simultaneously stimulated at wingbeat frequency, interference between stretch and neural activation results in variable work production. However, this interference disappears when the two activation timescales are close to each other resulting in entrainment to the neural drive. Matching time scales suggests an evolutionary path for smoothly transitioning to stretch-activated, asynchronous flight and back again.

## 1 Introduction

Muscles produce complex, dynamic forces that vary with length, activation, velocity, and prior strain trajectory. These dependencies of muscle force on state and history are nonlinear, making it difficult to predict their interaction during dynamic movement and to understand how new modes of force production have evolved to facilitate specialized locomotion.

Insect flight muscle is evolutionarily tuned for the incredibly high energetic demands of flapping flight (Fig. 1a). To achieve high-powered flight at ultrafast (100-1000 Hz) wingbeat frequencies, (*1, 2*), insects have evolved two types of flight muscle that reflect distinct ways of generating rhythmic oscillations in locomotion (Fig. 1b-c). Slow-flapping insects (*<* 100 Hz) rely on neurogenic activation of muscle which produces force synchronous with time-periodic nervous system signals (*3, 4*). However, fast-flapping orders of insects have evolved asynchronous flight muscle that is further activated by stretch, decoupling force production from the nervous system and enabling rapid wingbeats through self-excitation (Fig. 1b). Stretch-activation contributes to these insects’ ability to flap significantly faster than the 100 Hz barrier typically encountered by synchronous insects (*5*). While repeated evolutionary transitions between these flight modes have occurred, the minimal physiological properties needed to produce asynchronous force in a synchronous muscle remains unknown (*5*).

**Figure 1:**
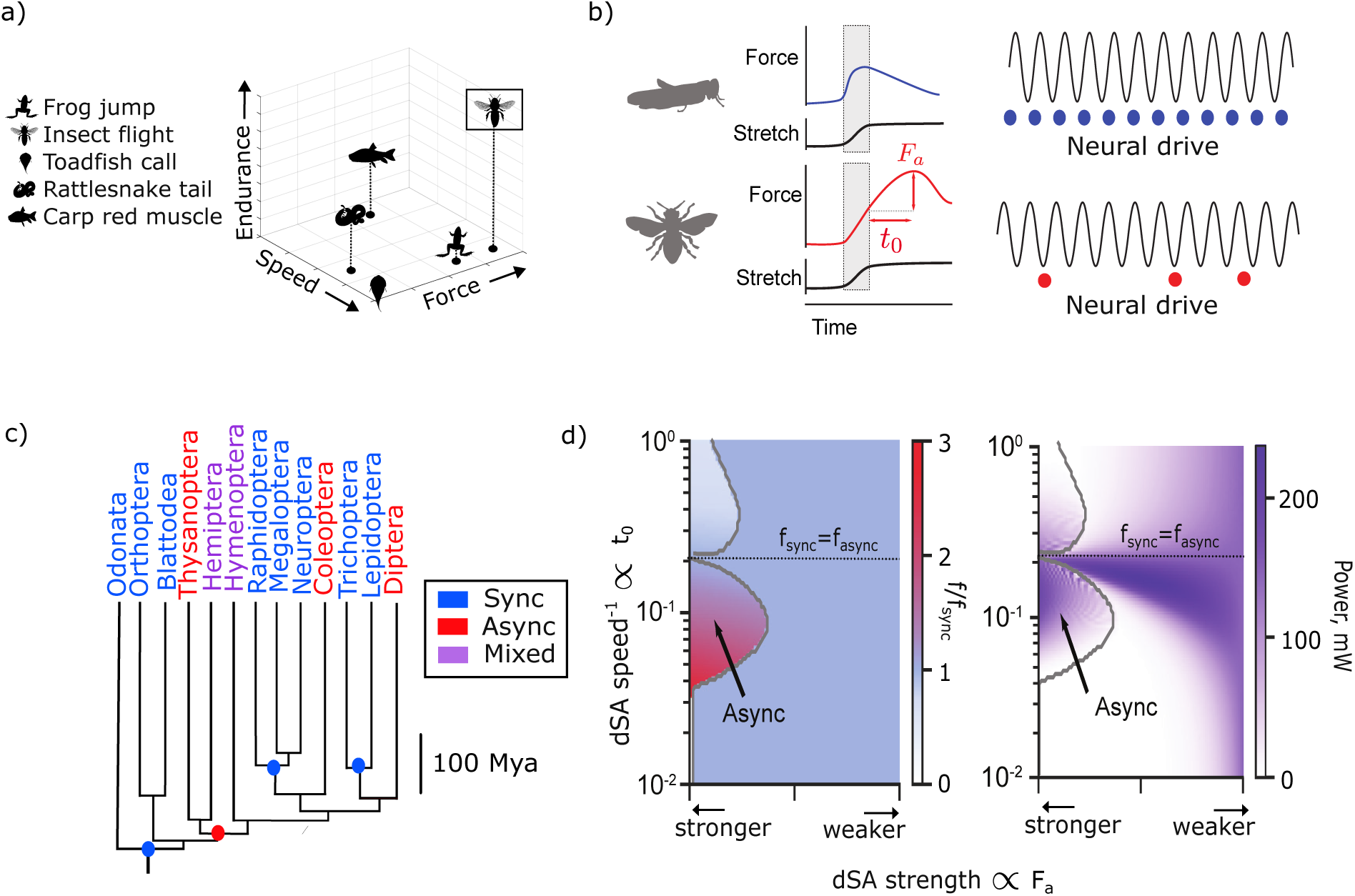
a). Skeletal muscles specialize over three main functional axes: speed, force, and endurance (*33*). Insect flight muscle must achieve high performance in all three categories. b). Delayed-stretch activation (dSA) decouples wingbeat frequency from the frequency of motor neuron activation in asynchronous insects. A characteristic timescale (*t_o_*) and magnitude (*F_a_*) typify the dSA response across insects (*5*) c). Pruned time-calibrated phylogeny of insect orders colored according to their flight muscle type. Red and blue dots denote likely transitions from sync-async and async-sync respectively (*5*). Further transitions in both directions have occurred within orders. d). Synchronous and asynchronous flight can be realized by a model with unified dynamics. Transitions between the flight modes depend on a combination of dSA speed and magnitude. When the frequencies of sync and async force production are matched, stable wingbeats are achieved that mix both modes (*5*). High power wingbeats can be achieved along a narrow bridge through parameter space which is defined by an entrainment boundary. b), c), d), e) modified from (*5*).

Ultrastructural and physiological evidence points towards a synchronous ancestral condition for flapping insects (*2, 6*). Ancestral state reconstruction supports one gain of asynchronous physiology at the order level and multiple subsequent reversions to synchrony (*5*) (Fig. 1c). However, inference of flight muscle type from histological and kinematic data limits the resolution of this reconstruction. Key differences in the molecular machinery among asynchronous orders leave open the possibility of multiple origins of asynchrony that converged on stretch-activation at the whole-muscle level (*7–9*). Furthermore, the existence of clades with both synchronous and asynchronous flapping species such as Hemiptera and Hymenoptera is highly suggestive of many more transitions in both directions within orders (*2, 6, 10*) (Fig. 1c). Since aerodynamics constrains smaller insects to flap their wings faster (*11, 12*), asynchronous flight muscle likely enabled the miniaturization and diversification of insects by unlocking ultrafast (100-1000 Hz) wingbeat frequencies (*13*). In addition, reversions from asynchrony back to synchrony have likely occurred at the order level in Lepidoptera, and almost certainly within orders like Hemiptera (*5*). These reversions may have enabled the evolutionary re-emergence of large body size (*14*) and precise wingstroke-to-wingstroke neural control (*15*) (Fig. 1b). Given its importance to evolutionary diversification of flapping flight frequencies, we sought to understand the minimal physiological properties that can induce asynchronous behavior and whether these properties can coexist with the neurogenic physiology of ordinary insect flight muscle.

Simple models of stretch-activated muscle hint at a potential core process underlying asynchronous wingbeats (*5, 16*). These models treat stretch-activation as a delayed active force rise in response to a step in strain, similar to the rapid stretch the muscle naturalistically would experience as it gets stretched by its antagonist. The characteristic force response to this strain stimulus exhibits a delayed rise and fall in force that drives the wingbeat, known as delayed stretch-activation (dSA) (Fig. 1b) (*2–4*). The rate of stretch-activated force rise is proportional to wingbeat frequency across asynchronous species, and is distinct from that of viscoelastic recoil (*17*). The amplitude of the dSA force response is typically two to threefold the tetanic force capacity of asynchronous muscles (*17, 18*). When two such muscles are antagonistically coupled to the insect thorax and wings, contraction of one muscle stretches the other and the overall system can self-excite to stable oscillations (*i.e.* limit cycles) (*3, 19*). The emergent frequency and amplitude of these oscillations are dictated by a combination of the dSA properties of the muscle as well as the resonant properties of the insect thorax and wings (*16, 20*). In asynchronous insects, neural activation modulates overall force production by adjusting intracellular calcium levels but is not the direct driver of individual wingbeats (*18, 21, 22*).

While synchronous and asynchronous flight appear to require different physiology, a recent unified model of flight muscle explained how such transitions between the flight modes could occur based on the same underlying dynamics (*5*) (Fig. 1d). The model assumes that flight muscle force is the weighted sum of a periodic synchronous component and an emergent asynchronous component operating at different timescales. When coupled to spring-wing mechanics, this hybrid sync-async dynamical system reproduces transitions between discrete flight modes by continuously changing either or both of just two physiological ‘knobs’ (dSA speed and magnitude) (Fig. 1d). Importantly smooth transitions with high-powered wingstrokes through the mixed regime only occur when the timescales of neurally activated and stretch activated forcing are close to each other through entrainment (Fig. 1d, right). Entrainment is a dynamical phenomenon that appears when a self-excited oscillation and an exogenous, time-periodic force co-occur in the same system. If the frequencies of these two oscillations are sufficiently close, the self-excited oscillation will spontaneously jump to the frequency of the exogenous force, generating a coherent single-frequency oscillation. While the model predicts the sufficiency of stretch activation for triggering a transition to asynchrony and the possibility of coexistence and entrainment with neurogenic oscillations, we do not yet know if insect flight muscle can actually realize these properties.

A gain-of-function experiment would be the gold standard to determine whether asynchronous behavior - the emergence of stable, positive work production from stretch-activation - can manifest from the layering of dSA on top of real synchronous muscle physiology. However, the precise molecular mechanisms of dSA are not known, and could differ between insect orders (*8, 9, 23*), rendering targeted genetic manipulation challenging. In the current work, we take the unconventional approach of giving stretch-activated properties to muscle using a technique we call the physiological dynamic clamp, inspired by dynamic clamp experiments in neuroscience (*24, 25*).

Akin to dynamic clamp experiments used to manipulate electrical coupling between isolated groups of neurons with analog electronics (*26–28*), we use a cyber-physical system to endow real muscle with a novel mode of force production. This approach builds upon previous closed-loop muscle physiology approaches that simulate embodied forces in real-time around a muscle (*29–32*) by simulating dSA with an in-silico model and adding it to the synchronous force produced by the muscle under neural activation. In doing so, we place a synchronous muscle in a virtual reality environment where it behaves as if it has dSA. This setup allows us to precisely control aspects of the synchronous and asynchronous components independently without affecting the dependence of muscle force on strain, strain rate, and activation. The physiological dynamic clamp provides an avenue to test how different modes of muscle force production combine without high-fidelity muscle models or genetic perturbations that may have off-target deleterious consequences for muscle function.

Here, we use the physiological dynamic clamp to give dSA to a normally synchronous moth flight muscle. We hypothesize that a synchronous muscle that has gained dSA should be able to realize stable asynchronous behavior when calcium-dependent force fluctuations are suppressed by inducing a near-tetanic state. This would suggest that the acquisition and loss of dSA is sufficient to drive transitions between synchronous and asynchronous flight, as has been hypothesized to have occurred over evolutionary time (*5*). Then, we manipulate the frequency of asynchronous force production to test the hypothesis that synchronous muscle can produce stable asynchronous power by entrainment of stretch-activated forces to a periodic neurogenic forcing. In doing so, we illuminate the conditions under which stretch and neural activation can coexist and the dynamical mechanism underlying their coexistence, providing a physiological basis for repeated evolutionary transitions in flight mode across insects.

## 2 Results

### 2.1 Modulating muscle properties in real time with the physiological dynamic clamp

We developed a novel virtual reality system to modulate muscle properties in real time called the physiological dynamic clamp. Briefly, the setup consists of an ergometer-mounted excised flight muscle that is coupled to a real time simulation (Fig. 2a). The simulation takes force and strain input, feeds them through a spring-wing model that captures the embodied (inertial, aerodynamic, and elastic) forces the muscle would interact with through the insect’s body, and outputs a length command back to the ergometer (Fig. 2b). In parallel, we send muscle strain into a simulation of dSA that computes stretch-activated forces based upon prescribed dSA rate and strength (Fig. 2c-e). We then add this stretch-activated force to the neurogenic force from the real muscle, and feed this into our spring-wing model. By changing parameters in the spring-wing and dSA models, we can give a real moth muscle with precisely tunable stretch-activated physiology. Using this setup, we were able to reproduce synchronous behavior in closed loop akin to traditional work loop experiments (Fig. 2f). We then layered dSA on top of the muscle’s neurogenic forces in a tetanic-like state mimicking the conditions present in asynchronous muscle (Fig. 2g), and during in-vivo-like activation patterns to understand how synchronous and asynchronoous forces interact in the same muscle (Fig. 2h). See Materials and Methods for a more detailed description of the setup and muscle preparation.

**Figure 2:**
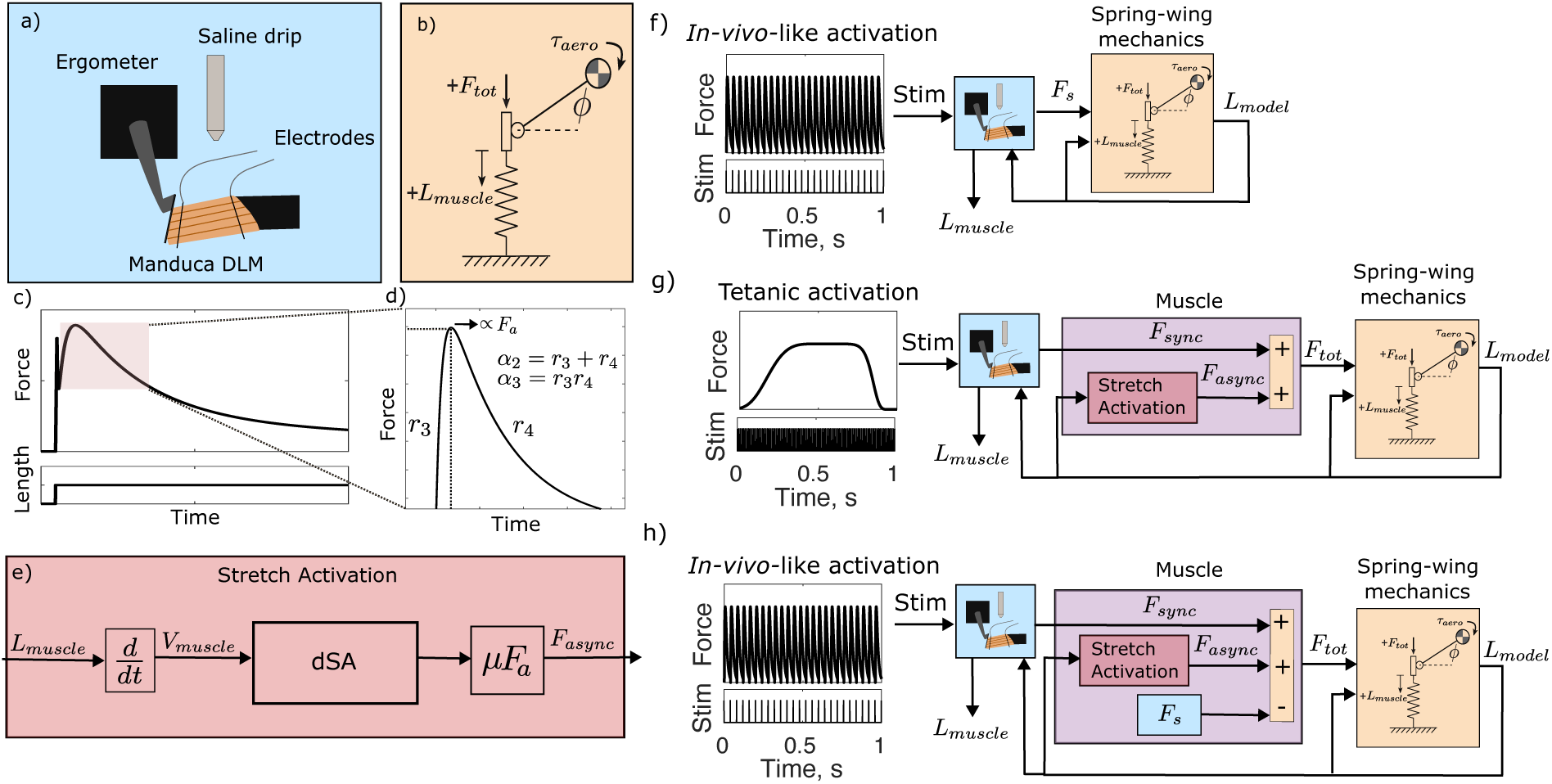
The physiological dynamic clamp for enhancing stretch activation in moth flight muscle. a) Schematic of *Manduca* flight muscle mounted to an ergometer. b). A spring-wing mechanical model takes muscle force and position input, computes virtual elastic and aerodynamic forces, and outputs a position command that is fed back to the ergometer and model for the next time step (*5*) The body mechanics module takes constant prescribed stiffness, damping, and inertial parameters that have previously been measured for *M. sexta* (*34, 35*). c) dSA is modeled as the muscle’s two-phase response to a step in strain. d) The response (Methods Eq. 2) is parameterized by a rate of force rise (*r*_3_) and a rate of force decay (*r*_4_) which can be reparameterized to *α*_2_ and *α*_3_, and a strength (*F_a_*) (*5*) e) A simulation of dSA (Methods Eq. 3) uses these parameters to implement a transfer function model that takes muscle position as an input and outputs a stretch-activated force. The transfer function is defined in the frequency domain with *s* denoting the Laplace variable. f). The closed-loop setup can be used without the physiological dynamic clamp to reproduce synchronous muscle behavior by feeding the neurogenic muscle force directly into the spring-wing model. g-h). The physiological dynamic clamp is used to layer dSA on top of neurogenic forces in real-time. In the first experiment (g) the flight muscle is held under tetanic electrical activation, mimicking conditions in most asynchronous muscles. by suppressing calcium fluctuations. In a second experiment (h), in-vivo-like neural stimulation is introduced to examine behavior of a muscle with strong synchronous and asynchronous physiology. A nominal synchronous force trajectory *F_s_* is measured prior to each experiment and subtracted in real time from the sum of synchronous and asynchronous forces to ensure length oscillation amplitude roughly matches *in-vivo* conditions (*36*).

### 2.2 Stable synchronous behavior in closed-loop in vitro virtual reality

We first tested our ability to elicit stable synchronous behavior in closed-loop from *Manduca* flight muscle in virtual reality. Prior workloop studies have shown that moth flight muscle necessarily produces positive mechanical work under conditions that replay in-vivo contraction strains and activation. However, we first needed to demonstrate that closing the loop between the muscle and biomechanical model of the springy insect exoskeleton and wings could produce stable oscillations. We stimulated the muscle periodically at *f_sync_* = 25 Hz, the *in vivo* frequency during flight, and observed the emergent length and force oscillations from its interactions with the spring-wing model (Fig. 2f). Stable length and force oscillations were consistently driven (Fig. 3a-b) at a frequency equal to *f_sync_*, indicating that the forcing from the real muscle, and not resonant self-oscillation of the spring-wing system, was determining the length oscillations. Because feeding back the prescribed length signal *L_model_* was necessary to ensure closed-loop stability, the length of the muscle measured from the ergometer *L_muscle_* is slightly time-lagged with respect to *L_model_* (Fig. 3a). Plotting the same measured force from the ergometer *F_s_* against each length trace, we see that a consistent workloop trajectory with the same frequency and amplitude results regardless of which length trace is used. Due to the time lag between *L_muscle_* and *L_model_*, work computed with *L_muscle_* is negative. This is a constraint of the virtual reality setup during synchronous activation - the delay between the two length signals is sufficient to push the muscle’s emergent phase of activation from a positive to negative work regime. Average work computed using *L_model_* was 1.85 ± 0.56 J/kg, within the range reported for *Manduca* (1.6-3.2 J/kg). Regardless, the virtual reality setup is able to reproduce similar amplitude and frequency length and force oscillations found in prior feedforward workloop studies (*37, 38*)

**Figure 3:**
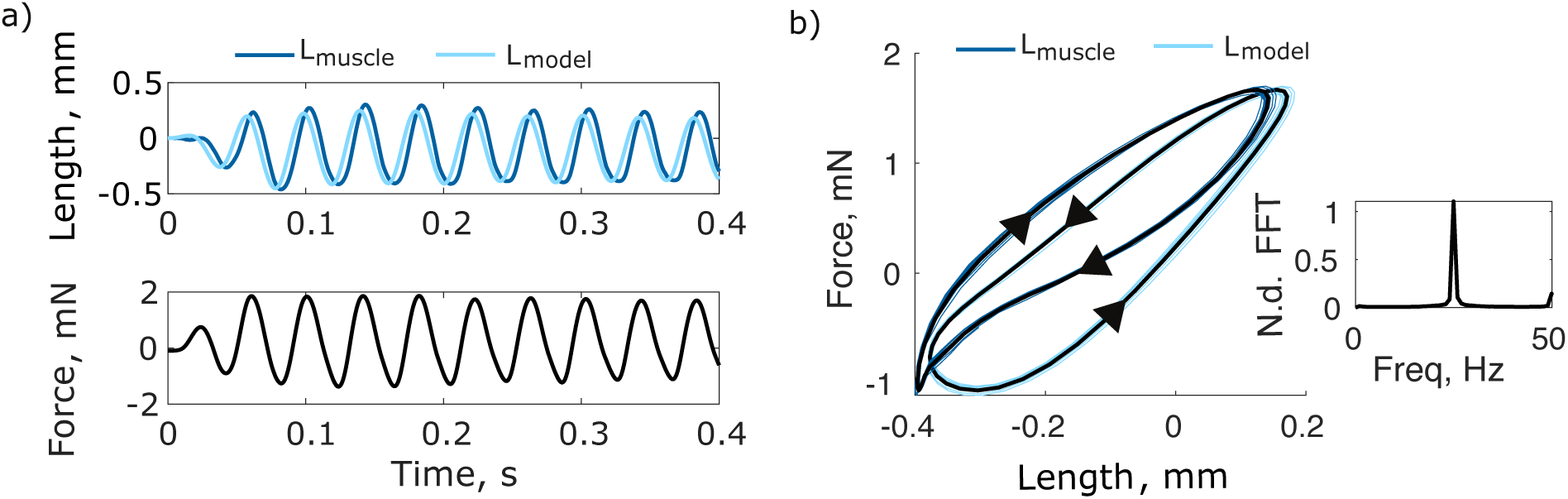
a). Synchronous, closed-loop oscillations from a *Manduca* flight muscle in virtual reality. Timeseries of the first few oscillations from one trial demonstrating stable length and force oscillations. *L_model_* is similar to *L_muscle_*, but slightly time-advanced. b). Workloop representations of the trial from which a) is plotted. The same force is plotted against each of *L_model_* and *L_muscle_*. Blue lines show all oscillations while black bolded line shows the mean trajectory. Inset shows the normalized FFT of the force, which has a single peak at *f_sync_* = 25 Hz.

### 2.3 Physiological clamping of stretch-activation produces asynchronous oscillations

Evolution of asynchronous muscle likely required an initially neurally activated muscle to have gained the capacity to produce stretch-activated work. Using the physiological dynamic clamp, we set out to characterize the mechanical (thorax-wing resonance) and physiological (stretch- and neural-activation) conditions under which stable, positive asynchronous work can be achieved in synchronous muscle. First, we attempted to elicit asynchronous behavior from synchronous muscle under tetanic activation. Tetanizing a synchronous muscle results in a plateau of force, which is maintained by high concentrations of calcium released from the sarcoplasmic reticulum (*4*). Most asynchronous insect flight muscle has very slow calcium handling which reduces the need for large volumes of sarcoplasmic reticulum and slows fluctuations due to neural activation (reducing *f_sync_*). During flight, the intracellular calcium concentrations inside asynchronous muscle are steady, producing a weak tetanus-like state. Calcium levels can still be modulated to control flight power (*21, 22*) but only over time scales much slower than the wingbeat frequency. The tetanized state of the synchronous flight muscle in the physiological dynamic clamp therefore mimics the slow calcium fluctuation conditions in asynchronous muscle during normal operation (*3*). We predicted that at near-tetanus, neurally-activated force fluctuations will be negligible, enabling positive asynchronous work production with a stable frequency and amplitude. This result would reject the alternative hypothesis that modeling of more complex stretch-activation dynamics or calcium-dependent modulation of stretch-activation (*21*) is required to manifest asynchronous behavior in an otherwise synchronous muscle.

We tetanized synchronous *M. sexta* muscle while adding simulated stretch-activated forces in closed-loop on top of its native synchronous forces with the physiological dynamic clamp (Fig. 2f). Our tetanic stimulation elicited average forces exceeding 1 N prior to the ramp-up of stable oscillations (Fig. 3a), indicating good muscle activation (*5*). We found that synchronous moth muscle produces stable, stretch-activated length and force oscillations under tetanus when virtual stretch activation is clamped in (Fig. 3b-c). The frequency of these oscillations was on average 20 Hz, far below the frequency of neural stimulation (100 Hz). However, this frequency is consistent with simulations of our asynchronous model - slightly faster than the resonant frequency of the simulated mechanics (*5*). Asynchronous wingbeat frequencies are set by a tug-of-war between the intrinsic rate of stretch-activation, *r*_3_, and the resonant frequency of the body and wings (*16*). When *r*_3_ is sufficiently fast with respect to the resonant frequency, asynchronous wingbeat frequencies can exceed resonance. Measurements from hawkmoths have demonstrated their wingbeat frequency during natural flapping is faster than resonance (*34*), and that their *r*_3_ is consistent with what would be expected from an asynchronous insect with their wingbeat frequency (*5*). Thus, emergent frequencies slightly faster than the resonant frequency in our experiments are consistent with previous models and are also comparable to wingbeat frequencies in large asynchronous hemipterans like *Lethocerus indicus* (*17*).

The stretch-activation length and force oscillations produce positive work along a stable periodic trajectory in force-displacement space (Fig. 3c). The muscle mass-specific work of these asynchronous oscillations was on average equal to 1.77 ± 0.31 J kg^−1^, within the range for *M. sexta* flight muscle’s synchronous work output (1.6-3.2 J kg^−1^,) (Fig. 3d) (*38–40*). Muscle mass specific power was 38.68 W/kg, slightly lower than the 40-80 W/kg previously reported for *Manduca* muscle, however this can be explained by the slightly lower frequency of asynchronous oscillations (18-20 Hz as opposed to 25 Hz during free flight). Thus, the muscle is capable of producing stretch-activated mechanical power equivalent to typical synchronous conditions. Furthermore, this power output was not diminished by nonlinear interactions between neurally-mediated and stretch-mediated force production. While *Manduca* does have a small degree of native dSA capacity, it is not strong enough to produce significant power compared to neurally-mediated forces. Our results show that if *Manduca sexta* flight muscle had stronger dSA with the same rate, it could produce stable asynchronous wingstrokes of sufficient power for flight at wingbeat frequencies close to what it uses for flight.

Contrary to the expectation of our model, which predicted oscillations of the same frequency across individuals as long as the model parameters remain unchanged, we observe highly variable frequency from preparation to preparation. We find a strong correlation between the active muscle stiffness (i.e. the slope of the work loop in Fig. 4c) and the emergent oscillation frequency (Fig. 4e) (*r*^2^ = 0.94). Thus, the oscillations that emerge in closed-loop are likely the result of additional stiffness in the real muscle on the ergometer, which elevates the emergent frequency beyond the expectation from the stiffness in our mechanics model. This result is perhaps unsurprising given that we are supra-maximally activating a synchronous muscle which stiffen much more than asynchronous muscles when neurally activated (*3*). This stiffening is further exacerbated by nearly tetanizing the muscle which is not typical for synchronous muscle in vivo. Regardless, we find that stretch-activation is sufficient for a synchronous muscle to generate positive stretch-activated work when neurally-activated force fluctuations are suppressed in tetanus.

**Figure 4:**
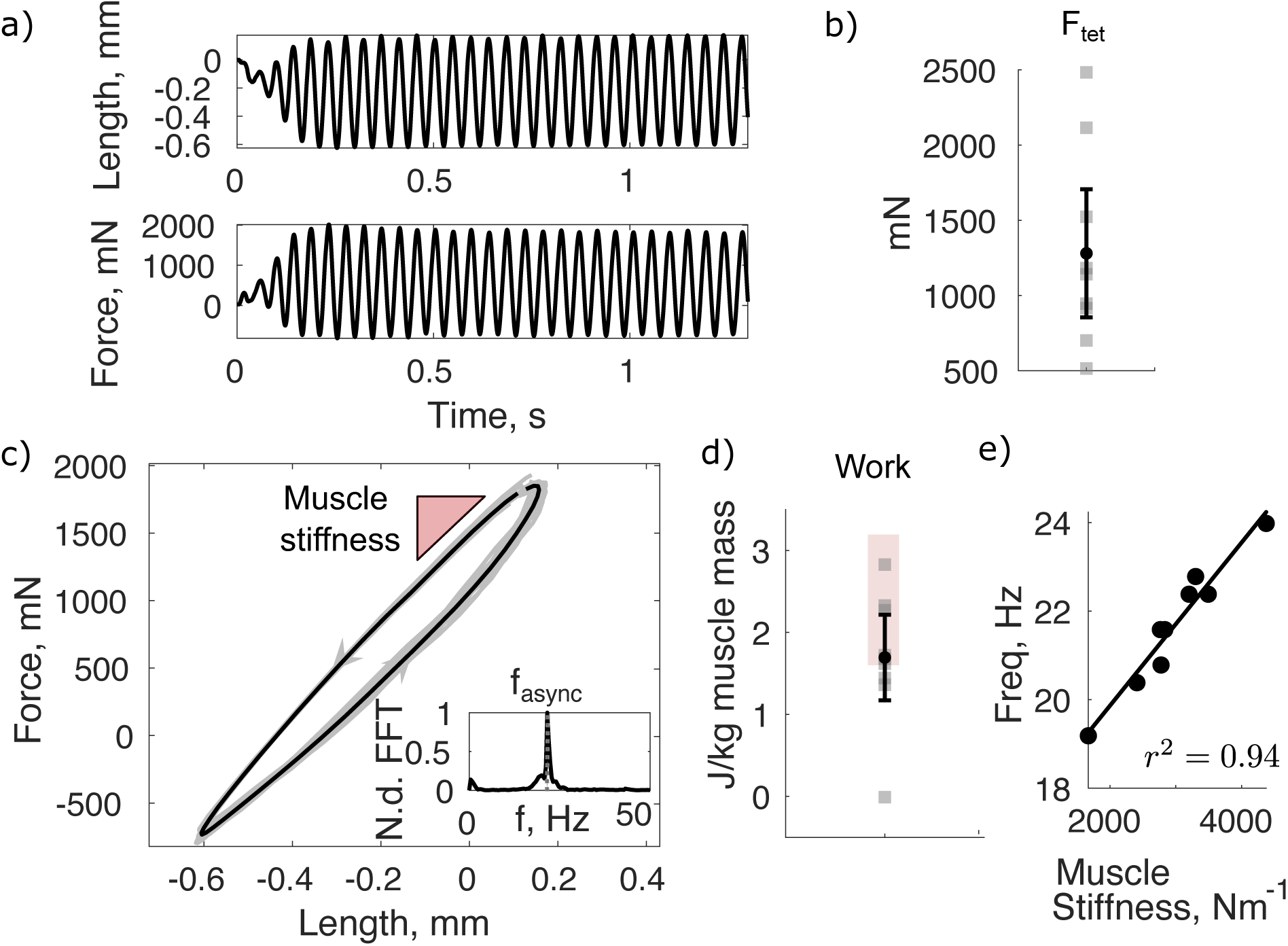
a) Example timeseries of measured length and combined sync-async force oscillations. b) Tetanic force *F_tet_* measured during the first 0.1s of stimulation was consistently above 1 N across preparations. c) The timeseries from a) plotted in force-length space. Time progresses in a counter-clockwise direction indicating positive overall work production. Inset non-dimensionalized FFT shows the dominant frequency of oscillation, corresponding to *f_async_*. Muscle stiffness can be approximated by the slope of the loop. d). Mass specific work across all individuals (n=10). Black dot shows the mean and bars show 95% CI of the mean. Red shading denotes range meausred from traditional open-loop work loop experiments. e). Emergent frequency *f_async_* is linearly related to the active stiffness of the muscle.

### 2.4 Asynchronous forces entrain to the nervous system under *in-vivo* activation

Next, we reintroduced realistic patterns of neural activation matching *in-vivo* conditions to the synchronous muscle. To explore how neural and stretch activation oscillation can co-exist in the same muscle, we tested whether periodic synchronous forces interfere with asynchronous forces as we manipulated the emergent asynchronous frequency. We hypothesized that the two modes of force production should combine as coupled oscillators with different frequencies, resulting in stable behavior over a classic entrainment boundary (*41, 42*). Alternatively, synchronous muscle’s myriad state- and history-dependent properties (*43–47*) may preclude stable asynchronous power production altogether, or destructive interference between flight modes modes may result in unstable power production under large sets of physiological or mechanical conditions.

We incorporated synchronous electrical stimulation mimicking *in-vivo* neural activation conditions by stimulating the muscle at 25 Hz while clamping in stretch-activated forces in closed-loop (Fig. 2h). The emergent hybrid oscillations are the result of processes with two different frequencies: the neurally driven fluctuations occurring at *f_sync_*, and stretch activation due to the physiological dynamic clamp at *f_async_*. While *f_sync_* is always set at 25 Hz, *f_async_* is not a directly set parameter. It depends on the interaction of the rate of stretch activation *r*_3_ and the resonant frequency of the spring-wing mechanics. Changing the effective stiffness in the model allows us to sweep through different values of *f_async_* by affecting the resonant frequency. Prior modeling showed that the forces due to both neural activation (synchronous) and stretch activation (asynchronous) act as oscillators with a state-dependent coupling between the two modes of force production (*5*). The state-dependent coupling places this system in a fundamentally different class from traditional phase coupled oscillators. While phase-coupled oscillators typically show entrainment, it is not clear that this must be a property for synchronous and asynchronous oscillators in muscle because of the coupling between exogenous (calcium) and intrinsic (stretch) activation. We hypothesized that the combined sync-async force will have both frequency signatures when *f_sync_* and *f_async_* are far apart (Fig. 5a,f). Interference between these frequencies will manifest in variable-amplitude oscillations and variable work production from cycle to cycle. However, when *f_sync_* and *f_async_* approach one another, we expect entrainment of asynchronous forces to the synchronous driving frequency manifesting in oscillations of a single frequency and stable amplitude (Fig. 5d). If this hypothesis is supported, then we can be confident that nonlinear interactions between the stretch and neural activation modes are not significant in the behavior of the hybrid system, and that moth flight muscle has the physiological potential to realize a continuous transition between synchronous and asynchronous oscillations.

**Figure 5:**
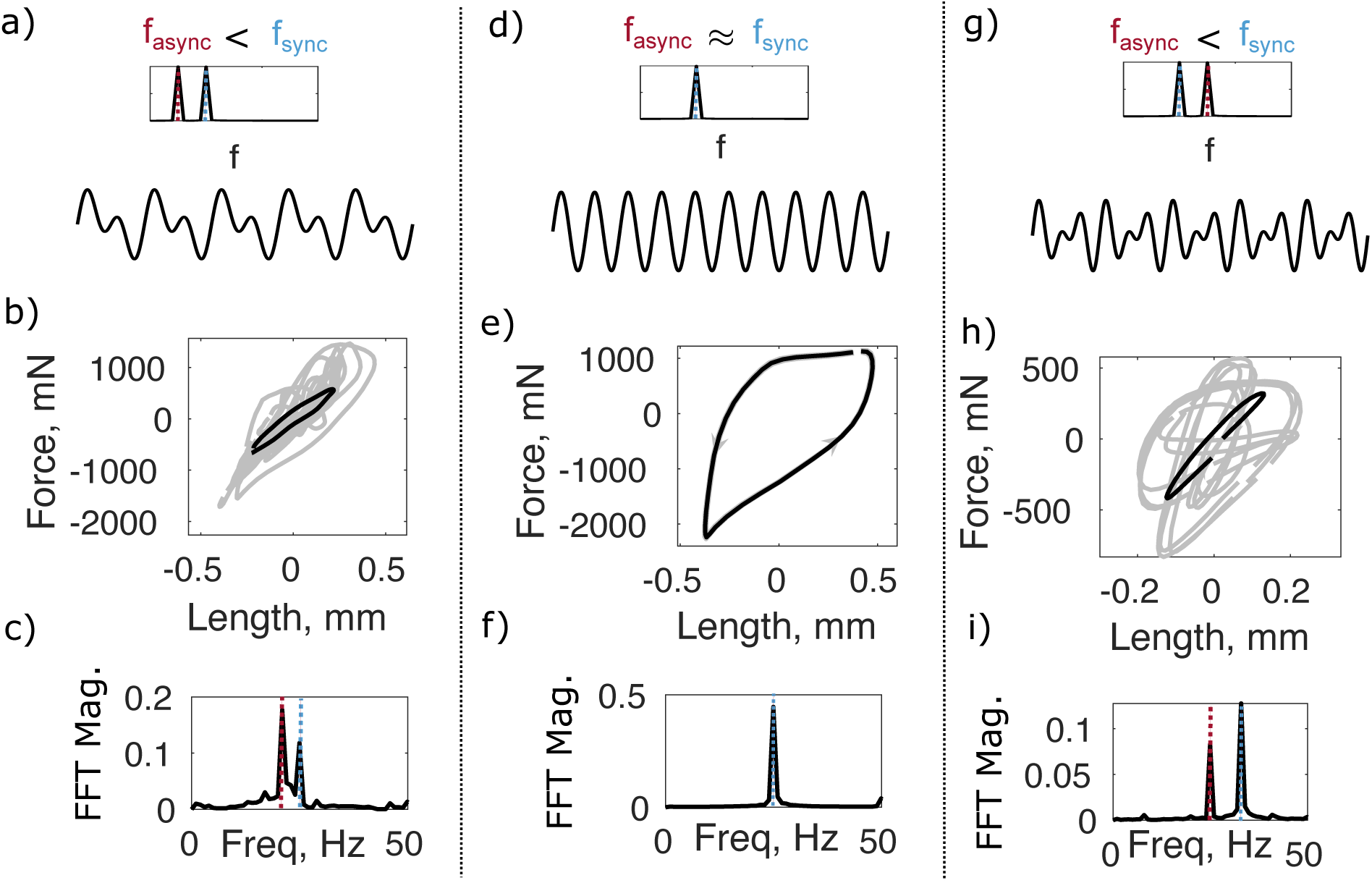
Predicted and measured frequency and amplitude of combined sync-async force for three different conditions: when *f_async_ > f_sync_* (a-c), *f_async_* ≈ *f_sync_* (d-f), and *f_async_ < f_sync_* (g-i). First row (a,d,g) shows the hypothesized relationship between *f_async_* − *f_sync_* and the frequency spectrum and amplitude of combined sync-async forces. Second row (b,e,h) shows measured work loops, and third row (c,f,i) shows measured the frequency spectrum associated with each set of *f_async_* and *f_sync_* conditions. Stable work loop trajectories with a constant frequency and amplitude only emerge when *f_sync_* ≈ *f_async_* (d-f). When *f_async_* and *f_sync_* are far apart (a-c, g-i), force amplitude fluctuates wildly and the frequency spectrum has signatures of both *f_async_* and *f_sync_*.

In accordance with our prediction, we observed highly variable work loop amplitudes when the neural stimulation frequency deviated by at least 3 Hz from the frequency excited by stretch activation (|*f_sync_* − *f_async_*| *>* 3 Hz) (Fig. 5a-c, g-i). The frequency spectra of these work loops contained two peaks, one corresponding exactly to *f_sync_* = 25 Hz and the other corresponding to *f_async_* (slightly above the resonant frequency) (Fig. 5c, i). Force and displacement amplitudes fluctuated from cycle to cycle, resulting in an average work loop that produced positive work equal to 1.75±0.24 J/kg (Fig. 5b, h). In a flying insect, this degree of cycle-to-cycle variation in muscle work would translate to highly variable lift production and unstable flight.

When *f_async_* was brought within ±3 Hz of *f_sync_*, we observed a single frequency peak equal to 25 Hz (Fig. 5f) (the neural driving frequency) and a work-loop trajectory without cycle-to-cycle amptlidue or frequency fluctuations (Fig. 5e). The consistent amplitude and frequency of these oscillations manifested in positive work output equal to 3.89±0.74 J/kg. Constant amplitude and frequency, and by extension low variation in cycle-to-cycle work production would result in consistent lift production inside the organism, consistent with the requirements for stable flight. A regime of stable oscillations driven by a combination of both activation modes would allow for evolutionary transitions in flight mode while maintaining strong flight capability throughout the transition.

Combining experiments across a wide range of *f_async_*, we see behavior consistent with entrainment of coupled oscillators (Fig. 6a-b). The width of the entrainment region was 7 Hz on average, demonstrating that entrainment occurs over a wide range of mechanical parameters (and hence a wide range of *f_async_*) (Fig. 6b). The width of the entrainment region corresponds to a frequency change caused by changing stiffness by ±50% or wing inertia by ±30% of the typical *M. sexta* values.

**Figure 6:**
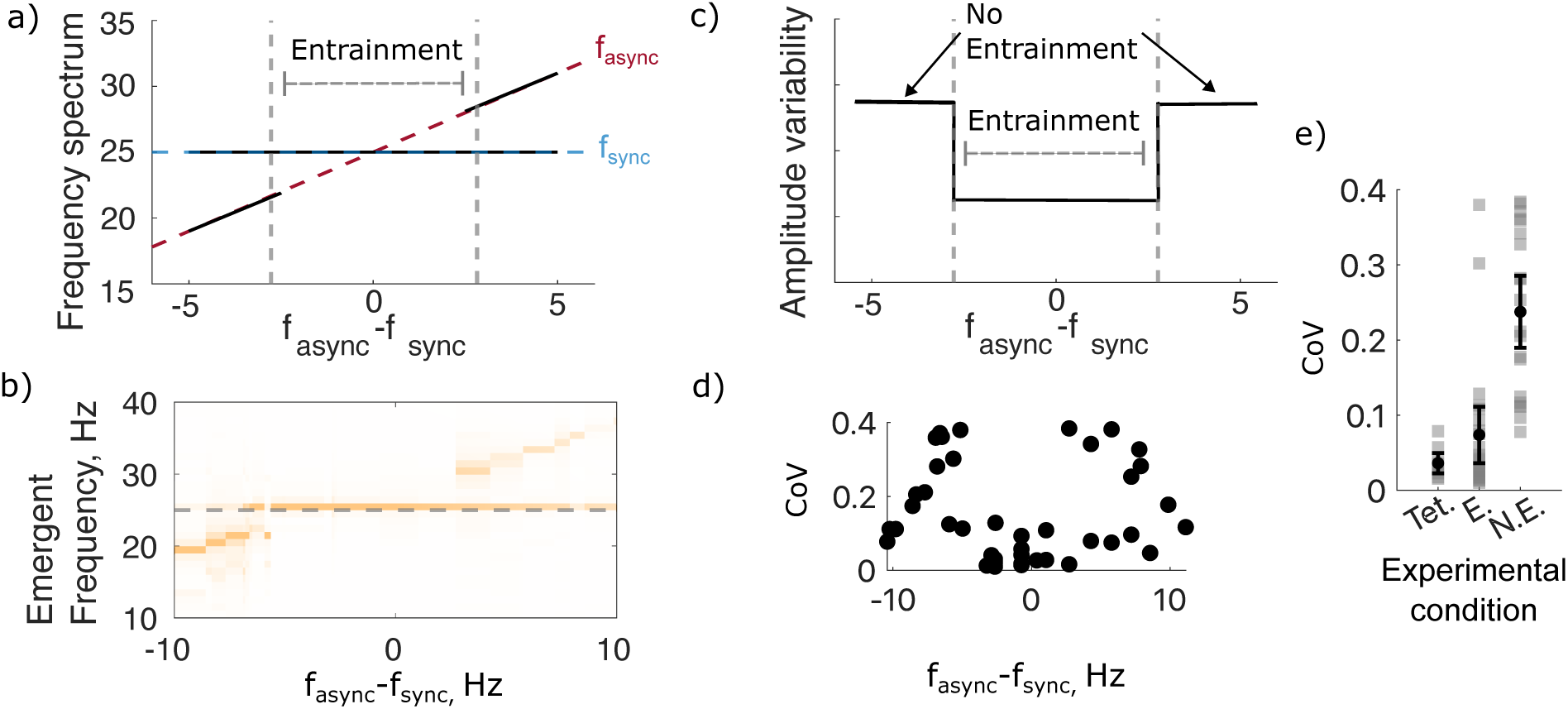
a) Prediction of entrainment boundary as a function of *f_async_* − *f_sync_*. When the two frequencies are far apart, the combined spectrum has two frequency components that correspond to the individual frequencies of *f_async_* and *f_sync_*. When the two frequencies are sufficiently close (in the entrainment region), the combined spectrum has only one frequency component since *f_async_* discretely jumps to *f_sync_*. b) The measured frequency spectra of trials from all individuals recapitulates the predicted entrainment pattern. Orange color intensity is proportional to FFT magnitude. c) Predicted amplitude variability as a function of *f_async_* − *f_sync_*. When the two frequencies are far apart, their interference results in a highly variable force amplitude. When the two frequencies are close (in the entrainment region), amplitude variability sharply decreases, reflecting the single frequency component of the combined sync-async force. d). Coefficient of variation (standard deviation divided by mean) of the emergent length oscillations recapitulates the prediction from entrainment, with low coefficient of variation when *f_sync_* and *f_async_* are close together. e) Coefficient of variation from trials conducted at tetanic (Tet.), entrained *in-vivo* (E.), and not-entrained *in-vivo* (N.E.) conditions. Tet. and E. trials have the same, lower variability than N.E. trials.

To quantify the consistency of the entrained oscillations between the different work loop conditions, we calculate the coefficient of variation across all cycles in a given trial (Fig. 6c-d). Computing this coefficient for both our tetanic and *in-vivo* activation trials, we found that the trials that exhibited frequency entrainment also demonstrated small amplitude fluctuations comparable to those under tetanus (Fig. 6e). In contrast, trials without frequency entrainment had large amplitude fluctuations as a result of interference between *f_sync_* and *f_async_*. Large amplitude fluctuations due to the presence of interfering frequencies would destabilize lift production in the organism which would be unfavorable for efficient flight.

## 3 Discussion

### 3.1 dSA is sufficient to realize asynchronous behavior

Using a novel physiological dynamic clamp approach, we demonstrate that a minimal model of delayed stretch activation is sufficient to drive asynchronous behavior in synchronous insect flight muscle (Fig. 4). The flight muscle of a hawkmoth can transition to a physiological regime necessary for asynchronous flight with the addition of dSA and produce sufficient frequencies and power to match what would be needed for flight (Figs. 5-6). Moreover, we did not need to change the time scale of stretch activation already present in *Manduca* flight muscle, we only needed to amplify its effect. Positive power is produced both under tetanic activation (Fig. 4c-d), and when paired with realistic neurogenic activation (Fig. 5e) demonstrating that the simple dSA model is robust to realistic synchronous force fluctuations. Strong, nonlinear interactions between dSA and neurally-activated dynamics would result in potentially unstable work production due to the emergence of higher frequency components. In this regard, our experiments show that any such interactions are unimportant to the steady-state behavior of the combined sync-async muscle system. The stable coexistence of both modes of activation in the same muscle supports the hypothesis that evolutionary transitions between the flight modes could have occurred continuously by passing back and forth through intermediate states of hybrid neural and stretch activation.

The two-parameter model of dSA (based on speed and magnitude) captures key features relevant for asynchrony, but is a simplified version of the actual processes that occur inside asynchronous myofibers. Classic characterizations of asynchronous muscle have employed a four-phase exponential response that includes two fast timescales of viscoelastic recoil as well as independent gains on each exponential (*48, 49*). Our results demonstrate that this level of complexity in the stretch response of asynchronous muscle is not strictly necessary to generate emergent stretch-activated positive power. Neither is explicit delayed shortening-deactivation (dSD), the decrease in force that occurs following rapid shortening and is the counterpart to dSA (*49*). While it does not include dSD, we note that the two-parameter model of dSA does respond to negative stretch, simulating the action of the antagonist muscle necessary to generate self-excited oscillations. Showing sufficiency of individual characteristics of stretch-activation for asynchronous behavior was not the goal of the aforementioned studies, and the full depth of stretch-activated properties are still likely important for asynchronous flight muscle function. dSD, for example is sometimes not observed in asynchronous muscle under certain ionic conditions, suggesting it may be an added specialization to fine-tune the stretch-response (*50, 51*). Thus, while a more detailed model of muscle stretch-response that incorporates other kinetic processes may allow for modifications of stretch-activation across insects, our results suggest it is not necessary for stable asynchronous power production.

While we do not explicitly test hypotheses related to molecular mechanisms of dSA, our results provide insight into which features of stretch-activation are needed to generate asynchronous behavior. dSA speed, or the rates of force rise and fall of the stretch response, is strongly related to wingbeat frequency (*17*) and thus is critical for tuning asynchronous power production. In the current work, we set these rates to match measured rates in *Manduca sexta* flight muscle, resulting in asynchronous wingbeat frequencies (20-24 Hz) close to hawkmoth free flight wingbeat frequency (25 Hz) (Figs. 4-5). The molecular determinants of dSA speed are not conclusively resolved (*52*). One candidate is the dissociation rate constant of myosin with respect to actin, a myosin ‘speed index’ which, in *Drosophila*, is among the fastest measured in insects. (*7*). Actomyosin dissociation as the rate-limiting step in stretch-activation is supported by the limited available comparative data across asynchronous insects, which suggest that the lifetime of the actomyosin bound state may determine maximum contractile speed (*52*). Evidence from other insects is strongly suggestive of multiple, potentially non-mutually exclusive mechanisms for stretch activation and asynchrony. Another promising candidate mechanism for dSA is the ratio of calcium-sensitive to stretch-sensitive myofilaments inside the sarcomere. Stretch-sensitive and calcium-sensitive isoforms of troponin have been discovered in asynchronous flight muscle, with their relative amounts correlated to asynchronous force production (*53, 54*). Tuning the relative abundance of these isoforms may allow for an increase or decrease in the strength of dSA over evolutionary time. Interestingly, the hawkmoth *Manduca sexta* also possesses stretch-sensitive troponin, suggesting a possible mechanism for its small, residual amount of dSA despite being synchronous (*55*). In the case of secondarily synchronous insects (synchronous insects with an asynchronous ancestor) like Lepidoptera, it is a reduction of dSA strength that allows neurally-mediated force production to dominate (*5*).

The thick filament myosin itself is stretch-sensitive, contributing to dSA in at least some insects. In *Drosophila*, a potential myosin-based mechanism for stretch-activation has been identified, likely driven by increased affinity of certain myosin isoforms for phosphate when stretched (*56, 57*). X-ray synchrotron experiments in flies, bees, and even frogs also suggest a myosin-based stretch sensing mechanism, which in insects may act in concert with the stretch-sensitive troponin C isoform (*58–60*). Titin-like molecules such as kettin, projectin, and flightin may also help transmit stretch across sarcomeres and directly to myosin heads (*52, 61, 62*).

Thus, there are likely many proteins responsible for setting both dSA speed and strength, and multiple molecular mechanisms that could give rise to similar stretch-activated behavior at higher levels of organization. Other sarcomeric properties seem to go hand-in-hand with known dSA mechanisms during the evolution of asynchrony, such as an increase in the ordering of the actomyosin lattice and reallocation of intracellular volume between organelles (*6, 63*). Our results support the idea that clade-specific mechanisms and regulatory proteins tune the shape of the dSA response, but may not be the core drivers of the delayed force rise itself. However, decoupling the common and ancestral components of asynchronous muscle behavior remains a promising area for further study.

### 3.2 Entrainment to the nervous system is consistent with an evolutionary bridge between synchrony and asynchrony

Stretch-activated and neurally-activated force production layer on top of one another in insect flight muscle as coupled oscillators, not linked by phase coupling but rather by the state dependence of stretch activation. In particular, our results are consistent with a model of a self-excited oscillation (stretch-activation) coupled to an external time-periodic forcing (neural-activation) (*41, 42*). When the frequencies of these two oscillations are far apart, the combined force exhibits signatures of both oscillation frequencies. When the frequencies are close, the combined force exhibits only the exogenously set synchronous frequency, irrespective of the asynchronous frequency. This is the hallmark of entrainment of self-excited oscillations to an external periodic forcing, in this case set by the nervous system (*19*), and has been observed in a robophysical simulation of mixed sync-async wingbeats (*5*). In the entrainment region, modulation of the neural driving frequency will more readily affect the self-excited dSA forcing, providing an avenue for neural control of stretch-activated wingbeats (*22*).

An important difference between our experiments and in-vivo dSA is that in-vivo, dSA properties are not fixed. Stretch-activated forces through dSA are themselves dependent on ambient calcium levels (*64*), which are set by synchronous neural input. However, this modulation is slow compared to individual wingbeats in most asynchronous insects, and is important for modulating flight power over longer timescales (*65*) and generating left-right asymmetries in power production for turns (*21*). In an insect operating in the entrainment region with a combination of synchronous and asynchronous muscle activation, synchronous activity may therefore modulate total muscle power by directly affecting stretch-activated forces, or by changing the frequency to which the stretch-activated forces entrain.

The discovery of the entrainment dynamics of asynchronous force production in synchronous muscle has implications for the evolutionary transitions between the two insect flight modes. The range of asynchronous frequencies over which entrainment occurred was at least 7 Hz in *Manduca* (Fig. 5b). Since asynchronous frequency variation is achieved by varying thorax and wing properties, this frequency range corresponds to a wide range of phenotypic variation (*5, 20*), providing a buffer of frequencies at which stable stretch-activated work can be achieved by a synchronous muscle.

Without entrainment, it is unlikely that a synchronous insect could evolve asynchronous flight by decreasing the strength of neurogenic force production and subsequently evolving stretch-activation. This would result in a flight-incapable intermediate phenotype. Evolution need not occur gradually, but entrainment provides a more likely scenario for a smooth transition between high-power intermediate, mixed-mode wingstrokes. An insect operating in the entrainment region can evolve stretch-activation without any required changes in neurogenic muscle physiology. Neurogenic force production may concomitantly or subsequently be tuned down to give rise to a fully asynchronous insect. Conversely, an asynchronous insect could evolve weaker dSA and stronger neurogenic force capacity while on the entrainment boundary. Once neurogenic forces dominate, it may then migrate off of the entrainment boundary. This is consistent with measurements from *Manduca* which place it off of the entrainment region with weak dSA (*5*). A large entrainment region may explain the frequency of repeated transitions in flight mode across the insect phylogeny (Fig. 1c) (*5*), providing a wide phenotypic space over which force-production modes can coexist without interference. Furthermore, entrainment supports repeated transitions in insect flight mode regardless of whether there was a single order-level origin of asynchrony or if multiple distinct types of asynchronous muscle have evolved across orders.

Beyond insect flight, stretch-activation is present to varying degrees in many other muscle types. The prevalence of dSA is likely underappreciated because it does not always manifest in standard *ex vivo* muscle conditions. Mammalian cardiac muscle exhibits moderate dSA (*66*) and certain mammalian skeletal muscles exhibit dSA that varies in magnitude depending on phosphate concentration (*56*). In the latter case the timescale of dSA has been shown to match the cycle frequency of mouse locomotion, and stretch activation is proposed as a potential adaptive mechanism to boost performance in fatigued states (*67*). The widespread nature of stretch-activation properties across muscle types, albeit likely the result of different molecular mechanisms, hints that stretch activation is generally beneficial in generating cyclic movements. Entrainment of stretch-activated forces to existing neural pacing in other muscle types may help explain its ubiquity, since stretch-activation properties need not be tuned precisely to the desired movement cycle frequency in order to be useful.

### 3.3 The physiological dynamic clamp

The physiological dynamic clamp is a technique that allows the experimenter to augment or modulate muscle physiological properties by coupling a real-time simulation to an isolated muscle preparation. This approach shares some similarities with other ‘closed-loop’ approaches in muscle biology, dating back to the seminal work of Machin and Pringle who used analog electronics to simulate virtual elastic, damping and inertial forces acting on asynchronous beetle flight muscle (*68*). More recent closed-loop muscle setups have been used to study how muscle forces interact with real-time dynamic loading, such as a frog muscle acting against an elastic or fluid substrate (*29, 31, 32, 69, 70*). The common thread through all prior closed-loop studies of muscle is the simulation of an external mechanical model with which a real isolated muscle interacts.

Our system starts with this closed-loop mechanical virtual reality environment. Instead of prescribing length or strain trajectories like classic work loops, we measure muscle force and feed it through the spring-wing resonant mechanics model. The physiological dynamic clamp innovates on the mechanical virtual reality environment by augmenting muscle physiology itself in closed-loop, not just a muscle’s interactions with external forces (Fig. 2f-h). This style of physiology experiment is perhaps most comparable to dynamic clamp experiments in neurophysiology (*24*). Neural dynamic clamp experiments use virtual gap junctions implemented with analog electronics to couple two physically separated clusters of cells, enabling the experimenter to precisely modulate intercellular electric coupling (*25, 27, 28*). Here, we implement a similar system in muscle physiology, using a dynamic clamp to simulate a specific mode of force production. Instead of modeling a gap junction with variable resistance, we model the physiological relationship of muscle strain and stretch-activated forces in asynchronous muscle (Fig. 2c-e). We then give this property to a synchronous muscle that lacks significant stretch-activation in a way that allows us to precisely manipulate each mode independently. In this way, we have developed a gain-of-function experiment for dSA despite not knowing its precise molecular mechanism, that can be applied in parallel with genetic or chemical physiological muscle manipulations.

Many kinds of muscle properties could be simulated and modulated using the physiological dynamic clamp. Other state and/or history-dependent properties of muscle such as work-dependent force modulation (*47*), length-dependent activation (*46*), residual force enhancement (*43*), or residual force depression (*71*) are ideal candidates for the closed-loop approach. The approach is limited only to virtual forces that can be adequately modeled as a function of muscle kinematics and prescribed model parameters. These virtual forces need not be expressible as the outputs of linear transfer functions, and thus can be used to study the effects of nonlinear phenomena on muscle function such as strain-stiffening elasticities (*72*) or muscle behavior in perturbative conditions (*44, 73*).

Our experiments also highlight the limitations of the closed-loop approach. Addressing hardware and signal transmission latency is critical, thus experimenters will benefit from using the fastest ergometry system available (*30*). Studying movements with a cycle frequency substantially slower than that of *Manduca* will make any unavoidable latencies less destabilizing to the overall closed-loop dynamics (Fig. 2e, Fig. 3) (*31*). In addition, certain properties of muscle that are easily controlled in open-loop are not easily controlled in closed-loop. For instance, phase of activation has strong effects on the work produced by most synchronous muscles (*37*), but cannot be easily prescribed in relation to the emergent length oscillations. These limitations may be surmountable by initiating closed-loop behavior with a transient open-loop length command and layering neural activation on top at a desired phase before transitioning the system to closed-loop.

Our work builds upon the rich history of cyber-physical systems in muscle physiology to give novel force-production capacity to an isolated muscle. We use this apparatus to show how two different types of oscillatory forces interact with one another inside the same muscle: time-periodic neurally-activated forces and self-excited stretch-activated forces. We are able to elicit stable positive asynchronous work production under different synchronous activation conditions, and show that a classic entrainment boundary allows for both modes of force production to co-exist without interference. In doing so, we demonstrate the utility of closed-loop approaches for answering questions in muscle physiology motivated by evolution.

## 4 Methods

### 4.1 In vitro virtual reality muscle physiology overview

We build upon previous *in-vitro* virtual reality systems for subjecting isolated muscles to realistic loading in real time (*29–32,69,70,74*). Our setup consists of a dual-mode ergometer (Aurora Scientific 305C), controlled by a Simulink Desktop Real-Time (SDRT) model at a sample rate of 1000 Hz (Fig. 2a). At every time step, length and force information is sent from the ergometer to the SDRT model, which then computes a length trajectory command that is sent back to the ergometer. Virtual reality simulates the embodied and environmental interaction forces a muscle would experience *in-vivo* around a real, isolated muscle, hence providing a virtual reality environment with real-time force and strain feedback for the muscle (*30*). The advantage of this approach is that it allows the systematic manipulation of mechanical parameters such as inertia, stiffness, and damping, while maintaining all of the biological complexity of real muscle-generated force production.

The SDRT model is comprised of two parts: a body mechanics module (Fig. 2b) and a muscle module (Fig. 2c-e). Simulating body mechanics in closed loop around an isolated muscle has previously been done using virtual reality, though only for much slower frog muscles (*29, 31, 32, 70, 74*). The body mechanics module takes muscle kinematic (i.e. position) input and computes the elastic, and aerodynamic forces that the muscle would experience based upon prescribed spring-mass-damper parameters. It then sums these virtual forces with the force from the muscle module, which by Newton’s second law, must equal the inertia multiplied by acceleration. These summed forces are then divided by the inertia (a constant parameter) and double integrated to yield a position command that is fed back to the ergometer and the mechanics model at the next time step (Fig. 2b).

The muscle module consists of the real force produced by the isolated synchronous flight muscle summed in real time with a simulation of dSA that takes muscle position as an input and outputs a stretch-activated force, depending on prescribed dSA rates and magnitude (Fig. 2c-e). This block implements a biophysical model of stretch-activation motivated by stretch-hold characterizations of isolated asynchronous muscle (*3, 5, 17*), and is described in detail below.

### 4.2 Muscle preparation and mounting

We isolate and mount the mesothoracic dorsolongitudinal flight muscles (DLM) of the hawkmoth *Manduca sexta* as has been done previously for *ex-vivo* studies of flight muscle mechanics (*37, 38*) (Fig. 2a). Briefly, moths were anaesthetized in a refrigerator before removal of the head, abdomen, legs, and prothorax. Compressed air was used to remove scales from the thorax exterior, and the thoracic ganglion was carefully snipped with dissection scissors to minimize residual nervous system activity to the flight muscles. The ergometer lever arm was glued with cyanoacrylate to the 2nd phragma, the posterior attachment point of the DLMs, via a pair of tungsten prongs. The scutum (anterior attachment point of the DLMs) was secured to a 3D-printed block clamped in place. Once the glue set, the thorax was set to its rest length, defined as the length at which the ergometer output read 0 N.

An incision on the dorsal exoskeletal surface was made and continued around the circumference of the thorax to separate its anterior and posterior hemispheres. Other musculature ventral to the DLMs was snipped to ensure the DLMs were the only muscle generating appreciable force during the prep. This dissection resulted in two, separate pieces of exoskeleton fixed on each side and connected only by the two DLM muscles. Once dissected, a temperature-controlled saline drip was turned on which supplied the muscle with a moth Ringer’s solution. A pair of tungsten-tipped silver wire electrodes were inserted through the residual dorsal cuticle into each DLM. Electrodes were placed towards the insertion and origin points of each muscle and pierced through all subunits to maximize neural activation of the muscle. Immediately prior to beginning an experiment, we raised the temperature of the saline drip to 34 degrees C, close to the internal temperature of freely-flying hawkmoths (*38*).

### 4.3 Simulink Desktop Real Time Model

We constructed a SDRT model that simulates forces that act upon the isolated moth muscle in real time using Simulink 2020b. In particular, we consider elastic forces generated by the deformation of the moths exoskeleton, inertial forces from the acceleration and deceleration of the wing mass, and aerodynamic forces from the generation of lift around the wing (Fig. 2b). These are the main forces experienced by moths in flight, and dominate unmodeled forces such as internal thoracic damping (*34, 35, 75*).

In any closed-loop motor control setup, time delays due to signal latency or hardware latency can be major problems, resulting in unstable resonant oscillations of the motor (*30*). In our system, the most salient source of delay is the ergometer itself. Upon receiving a length command, the PID module inside of the ergometer’s control box is responsible for ensuring the arm achieves the desired position. When custom tuned for maximum response time, our ergometer can accurately respond to a position command within 2 ms unloaded. This latency increases when a complex load is applied to the arm. Since the *in-vivo* wingbeat period of *M. sexta* is 40 ms (*15*), a latency of 1-2 ms is significant and destabilizes the closed-loop system, causing uncontrolled resonant oscillations. In previous muscle virtual reality experiments, this has not been an issue since the cycle period of motion was much larger with respect to the maximum latency (*31*).

To circumvent this problem, we feed back the prescribed motor position instead of the measured motor position to our muscle and mechanics model (Fig. 2f-h). Doing so ensures the model always experiences a signal latency of 0 ms and enables stable closed-loop behavior. However, this has the consequence of time-shifting the measured length oscillations from the prescribed length oscillations (output by our model) by approximately the ergometer latency. Our results are qualitatively unchanged regardless of which length trajectory is used to plot data and compute work output. To provide the most accurate reflection of the dynamic conditions actually experienced by the muscle we analyze the measured length signal as opposed to the time-advanced, prescribed length signal, but discuss both.

### 4.4 Spring-wing mechanics model

We modeled the body mechanics and aerodynamics of a flying hawkmoth by implementing a ‘spring-wing’ dynamical equation (*5, 34, 35, 76*) for the insect’s wing angle in closed-loop (Fig. 2b).

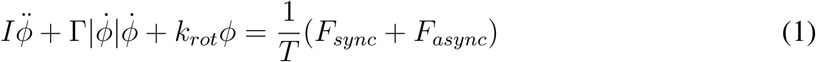

In this equation, *I* is the wing inertia, Γ is an aerodynamic force coefficient, *k_rot_* is the rotational stiffness of the wing hinge, and *T* is the transmission ratio which converts muscle force into torque about the wing hinge. *F_sync_* is the measured muscle force and *F_async_* is the simulated asynchronous force. All mechanical parameters were set to match *in-vivo* conditions for *M. sexta* (*5, 34*). Γ is a simple translational quasi-steady blade element model of aerodynamics defined as:

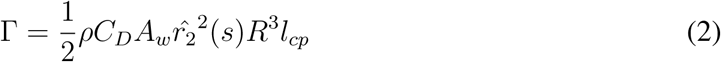

In the above equation, *ρ* is air density, *C_D_* is the empirically determined coefficient of drag, *A_w_* is the wing area, *r̂*_2_(*s*) is the non-dimensional second moment of wing shape, *R* is wing length, and *l_cp_* is the non-dimensional location of the center of pressure (*77*). More complex models of aerodynamics that incorporate wing pitching or other aerodynamic phenomena are unlikely to affect the emergent wingbeat frequencies. In principle this is because the resulting wingbeat frequency is determined primarily by the mechanical resonance of the system and the muscle physiological timescale. In practice, this robustness to aerodynamics is evidenced in both dynamically scaled and at-scale robophysical models.

### 4.5 dSA model

We use a recent dynamical model of stretch-activated force production to simulate asynchronous forces in real-time (*5, 78*). The model is based upon classic stretch-hold characterizations of asynchronous muscle. When an asynchronous muscle experiences a step in strain, it exhibits a four-phase force response well-described by the following functional form (Fig. 2c-d):

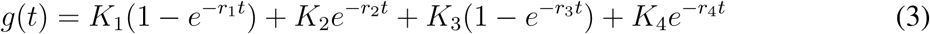

As in prior work (*5,16,78*), we simplify this four-phase response to just two phases, ignoring the first two exponential processes which correspond to the very fast viscoelastic recoil of the muscle following stretch. The latter two exponential processes comprise the much slower dSA force that actually drives the wingbeat. We additionally assume the coefficients (K’s) on each exponential process are equal to unity, resulting in a two-parameter description of dSA (Fig. 2d) (*5*).

We then convert this two-parameter dSA model into a differential equation that describes the dynamics of asynchronous force production (see (*78*) for a complete derivation of this procedure), where *α*_2_ = *r*_3_ + *r*_4_ and *α*_3_ = *r*_3_*r*_4_:

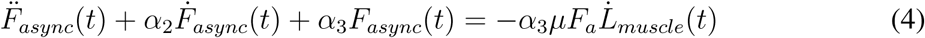

This equation can be implemented in closed loop as a transfer function that takes in *L̇_muscle_* (the time derivative of muscle position) at each timestep and computes *F_async_*, given a few prescribed parameters (Fig. 2e):

The two rates *r*_3_ and *r*_4_ have been measured for a variety of asynchronous insects and vary with wingbeat frequency (*17*). Thus, the frequency of asynchronous forces produced by the model will depend on these rates. Despite being a synchronous insect, *Manduca sexta* has weak dSA in its flight muscles, but with a timescale that is appropriately tuned for its wingbeat frequency of ≈ 25 Hz (*5*). Thus, we used the dSA rates measured in *Manduca* (*r*_3_ = 36 s^−1^, *r*_4_ = 22 s^−1^) to ensure our simulated asynchronous dynamics are roughly scaled to *in-vivo* relevant conditions. These rates primarily dictate the rate of asynchronous force production (i.e. frequency).

The strength of asynchronous force production is set by *µ* and *F_a_*. *F_a_* is taken from experiments to be the magnitude of the measured dSA force response (Fig. 2d) (*5*). However *F_a_* in *Manduca* is far too low to drive meaningful force production *in-vivo*. This low *F_a_* is the reason why *Manduca* remains synchronous despite having flight muscle with some dSA (*5*). To augment *Manduca*’s intrinsic dSA, we define a parameter *µ* that scales the strength of the dSA response, acting as a gain on the asynchronous force amplitude (Fig. 2e). We tune *µ* such that the emergent asynchronous forces drive length oscillations in the motor that are comparable to realistic muscle strain trajectories *in-vivo* (*36*). The constant value of *µ* used in all of our experiments was *µ* = 0.002.

### 4.6 Experimental protocol

We performed two sets of experiments with our closed-loop muscle physiology system, under two different activation conditions: tetanic activation (Fig 2f), and periodic activation matching *in-vivo* wingbeat frequency (Fig 2g-h). We endeavored to perform both experiments on each animal used, although usable data was not always obtained from both experiments. Tetanizing synchronous insect flight muscle for long periods of time is an extreme activation condition that we found to cause rapid deterioration of force production. Thus, we always conducted the tetanic trial last for each animal.

### 4.7 Synchronous closed-loop test

Before either of our two main experiments, we performed a test of synchronous closed-loop behavior. The muscle was coupled only to the spring-wing model without any simulated dSA and stimulated electrically at 25 Hz. The measurement of the synchronous closed-loop force response *F_sync_* served as a metric of preparation health and consistency. Preparations with low synchronous force production would yield an *F_sync_* dominated by the thorax-wing resonant frequency, since there was not enough synchronous force to overcome the motor’s tendency to resonate in closed-loop after a perturbation. Only preparations with an *F_sync_* that had a frequency of 25 Hz were continued.

#### 4.7.1 Tetanic activation condition experiments

In these experiments, we tetanized the muscle by stimulating it electrically with biphasic voltage pulses at 100 Hz (Fig. 2f). Stimulus strength was set to a value that maximized twitch force, usually around 6V. Doing so in an isometric muscle results in a fused tetanus (*5*). Tetanizing the muscle simulates the conditions under which asynchronous insects generate stretch-activated muscle force. In fast asynchronous muscle such as that of a bumblebee, the muscle is effectively tetanized (stimulated neurally at a high frequency), but force production is dominated by the stretch-activated mode (*79*), not the neurally-activated mode. We could not perform an open-loop tetanus as a control since doing so would damage the muscle. Instead we took the maximum force *F_tet_* as the force reached during the first 0.1 s of stimulation in closed-loop, before oscillations had time to ramp up. In these experiments, we held all body mechanical parameters constant (i.e. inertia, stiffness, damping, and dSA rates), and measured work, frequency, and amplitude from *F_tot_* and *L_muscle_*.

#### 4.7.2 *In-vivo* activation condition experiments

In these experiments, we re-introduced realistic activation conditions to the muscle by stimulating it at 25 Hz - the native frequency of stimulation during free flight (*36*) (Fig. 2g-h). Doing so in an isometric muscle results in a train of twitches that match the stimulation frequency of 25 Hz. However, simply adding the neurally-activated forces to the simulated stretch-activated forces in closed-loop presents some issues. If we approximately match the strength of the stretch-activated forces to the strength of native synchronous force production, the total force produced by the muscle will be larger (up to double, in the case of perfect constructive interference) than realistic forces produced in flight. This is a problem since we wish to approximately match the length oscillation amplitude of the hybrid system to realistic length oscillations during flight.

In an effort to reduce this effect, we subtracted the purely synchronous force response *F_s_* (Fig. 2e) from the total hybrid synch-asynch force in real time (Fig. 2h). This pre-measured force was the force produced by the muscle in closed-loop without any simulated dSA, while being activated at 25 Hz (Fig. 2g). Doing so reduces the amplitude of the purely synchronous forces enough that the combined hybrid muscle produces force with an amplitude that is comparable to that during realistic flight, while leaving interactions between the force production modes unaffected. We measured *F_s_* first in each animal, and used the same *F_s_* for the rest of the *in-vivo* experiments.

We manipulated the asynchronous frequency, *f_async_*, while leaving the synchronous frequency, *f_sync_* (equal to the 25 Hz stimulation), constant by changing the value of the stiffness *k_rot_* in our simulated mechanics mode (Fig. 2b). *f_async_ in-vivo* is emergent from interactions between the dynamics of dSA and the resonant mechanics of the thorax-wing system (*5, 20*). Thus, altering the resonant properties (i.e. stiffness or inertia) of the body mechanics model achieves the same effect on *f_async_* as changing the dSA rate *r*_3_. By varying the modeled undamped spring-wing resonant frequency 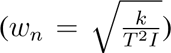, we induce a systematic, concomitant change in *f_async_* (*5,78*) without affecting *f_sync_*. In doing so, we can examine the extent to which the two modes of force production combine nonlinearly over a range of *f_async_*. Note that *f_async_* is usually substantially larger than the resonant frequency, but always changes in proportion to the resonant frequency (*5*).

### 4.8 Data analysis

Raw length and force data from the ergometer was read out from the PCIe card into MATLAB for processing. We considered the length of the muscle to be the prescribed length signal (i.e. not the time-delayed measured length signal). As described above, for stability reasons we used the prescribed length signal as our length feedback as opposed to the measured length from the motor. However, all plots and analyses in Figs 4-6 were done with the measured length signal to accurately capture the dynamic conditions experienced by the muscle. We analyzed the total force generated by the muscle block, *F_tot_*, which is the sum of the measured force and the simulated asynchronous force at each time point. Data remained unfiltered for all analyses. We computed frequency content by taking the fast fourier transform (FFT) of the measured length signal, though the frequency content of all three signals (measured length, prescribed length, and total force) were similar to one another.

